# A Bispecific Antibody Targeting RBD and S2 Potently Neutralizes SARS-CoV-2 Omicron and Other Variants of Concern

**DOI:** 10.1101/2022.05.11.491588

**Authors:** Mengqi Yuan, Xiangyu Chen, Yanzhi Zhu, Xiaoqing Dong, Yan Liu, Zhaohui Qian, Lilin Ye, Pinghuang Liu

## Abstract

Emerging severe acute respiratory syndrome coronavirus type 2 (SARS-CoV-2) variants, especially the Omicron variant, have impaired the efficacy of existing vaccines and most therapeutic antibodies, highlighting the need for additional antibody-based tools that can efficiently neutralize emerging SARS-CoV-2 variants. The use of a “single” agent to simultaneously target multiple distinct epitopes on the spike is desirable to overcome the neutralizing escape of SARS-CoV-2 variants. Herein, we generated a human-derived IgG-like bispecific antibody (bsAb), Bi-Nab_35B5-47D10_, which successfully retained the specificity and simultaneously bound to the two distinct epitopes on RBD and S2. Bi-Nab_35B5-47D10_ showed improved spike binding breadth among wild-type (WT) SARS-CoV-2, variants of concern (VOCs) and variants being monitored (VBMs) compared with its parental mAbs. Furthermore, pseudotyped virus neutralization demonstrated that Bi-Nab_35B5-47D10_ can efficiently neutralize VBMs including Alpha (B.1.1.7), Beta (B.1.351) and Kappa (B.1.617.1) and VOCs including Delta (B.1.617.2), Omicron BA.1 and Omicron BA.2. Crucially, Bi-Nab_35B5-47D10_ substantially improved neutralizing activity against Omicron BA.1 (IC_50_= 27.3 ng/mL) and Omicron BA.2 (IC_50_= 121.1 ng/mL) compared with their parental mAbs. Therefore, Bi-Nab_35B5-47D10_ represents a potential effective countermeasure against SARS-CoV-2 Omicron and other variants of concern.

**Importance:** The new highly contagious SARS-CoV-2 Omicron variant caused substantial breakthrough infections and has become the dominant strain in countries across the world. Omicron variants usually bear high mutations in the spike protein and exhibit considerable escape of most potent neutralization monoclonal antibodies and reduced efficacy of current COVID-19 vaccines. The development of neutralizing antibodies with potent efficacy against the Omicron variant is still an urgent priority. Here, we generated a bsAb, Bi-Nab_35B5-47D10,_ that simultaneously targets SARS-CoV-2 RBD and S2 and improved neutralizing potency and breadth against SARS-CoV-2 WT and the tested variants compared with their parental antibodies. Notably, Bi-Nab_35B5-47D10_ has more potent neutralizing activity against the VOC Omicron pseudotyped virus. Therefore, Bi-Nab_35B5-47D10_ is a feasible and potentially effective strategy to treat and prevent COVID-19.

## Introduction

Owing to the continuous SARS-CoV-2 evolution caused by mutations and recombination, numerous genetically distinct SARS-CoV-2 lineages have emerged, and five major variants, including alpha (B.1.1.7), beta (B.1.351), gamma (P.1), delta (B.1.617.2), and the newly identified micron (B.1.1.1.529 and BA), have been designed sequentially based on criteria such as various transmissibility and the ability to escape immunity[1-6]. Most mutations of SARS-CoV-2 concern variants primarily clustered on the NTD and RBD, two regions of spike S1, the two major domains being targeted by neutralizing antibodies[5, 7-10]. Many of these mutations in S1 have been previously reported to undermine the effectiveness of COVID-19 vaccines and therapeutic neutralizing antibodies[11, 12]. The recently identified SARS-CoV-2 Omicron variant of concern (VOC) worsened the situation[13]. The Omicron BA.1 variant harbors an unusually high number of mutations and rapid spread capacity. Omicron BA.1 has over 30 mutations in the viral spike protein, including 15 mutations in RBD[14]. The Omicron variant has rapidly replaced the previously dominant Delta variant and has become the dominant circulating strain in many countries across the world since it was first reported in November 2021 in Botswana and South Africa[15, 16]. The substantial mutations of the Omicron variant make it completely or partially resistant to neutralization by most potent monoclonal antibodies against other VOCs and result in significantly reduced efficacy of existing COVID-19 vaccines[17-20]. Therefore, it is still a top priority to develop highly potent and broadly monospecific or multispecific neutralizing mAbs targeting SARS-CoV-2 heavily mutated variants, such as the Omicron variant[21].

Monoclonal antibody cocktails or bispecific antibodies are an effective strategy to counter the escape of highly mutated SARS-CoV-2 variants[22]. The bsAb strategy has been successfully applied to the treatment of cancer and inflammatory disorders and viral infectious diseases[23-26]. Bispecific antibodies are advantageous over antibody cocktails given the complicated formulation and cost of antibody cocktail strategies[27-30]. The SARS-CoV-2 spike ectodomain is segregated into two units, termed S1 and S2. The S1 subunit of SARS-CoV-2 containing NTD and C-terminal RBD is responsible for cellular binding and is targeted by most neutralizing antibodies, whereas the S2 subunit is relatively more conserved and mediates membrane fusion[31, 32]. The S2 region, harboring neutralizing epitopes, is an alternative conserved target on the spike[33]. To avail the cellular binding and fusion of two key steps of SARS-CoV-2 entry, we generated a human-derived, IgG1-like bispecific antibody Bi-Nab_35B5-47D10_ based on two human neutralizing antibodies, 35B5 and 47D10, which target the spike RBD and S2 region, respectively. Human mAb 35B5 is a potent human monoclonal antibody panneutralizing against WT SARS-CoV-2 and VOCs, whereas 47D10 is an anti-S2 human neutralizing antibody with crossing activity against several beta coronaviruses. Our results show that Bi-Nab_35B5-47D10_ neutralizing activities successfully maintained parental specificity and simultaneously targeted two epitopes of the RBD and S2 region, and the two arms of this bispecific antibody potently neutralized various circulating SARS-CoV-2 variants. Notably, it has improved neutralization activity against neutralizing relatively resistant SARS-CoV-2 Delta and Omicron BA.1 variants. These results indicate the potential development of therapeutic strategies of Bi-Nab_35B5-47D10_ against SARS-CoV-2 Omicron and other variants.

## Results

### Design and expression of bispecific antibodies

To obtain functional bsAbs, we designed and generated bsAbs containing tandem single-chain variable fragment (ScFv) domains of two potent neutralizing mAbs (35B5 and 47D10) with G4S linker separation based on structural information and computational simulations of spike trimers as previously described [34]. We selected 35B5 and 47D10 as two neutralizing antibodies given that their epitopes are located in different regions of the spike and lack an unfavorable steric clash. The 35B5 mAb is a new class RBD-targeting potent neutralizing human antibody through a distinctive spike glycan-displacement mechanism, and its epitope is invariant among SARS-CoV-2 wild-type and circulating variants [35, 36]. The 47D10 mAb, isolated from WT SARS-CoV-2-infected patients’ memory B cells, was identified as a SARS-CoV-2 S2-specific mAb (Fig. S1A and B). Based on 35B5 and 47D10, we designed two molecular topologies of bsAbs: 35B5_VL-VH_ → G4S linker → 47D10_VL-VH_ (Fig. 1A) and 47D10_VL-VH_ → G4S linker → 35B5_VL-VH_ (Fig. B). To prevent possible steric resistance caused by the SARS-CoV-2 spike trimer, the empirical choice of 5X G_4_S linkers was made. To maintain the antibody Fc-mediated activities, the individual 35B5 and 47D10 ScFvs were then connected using 5X G_4_S linkers and fused to the IgG1 Fc to generate the IgG-like bsAbs molecule (Fig. 1C). Two bsAbs, Bi-Nab_47D10-35B5_ and Bi-Nab_35B5-47D10,_ were produced by using the ExpiCHO^TM^ Expression System and purified by affinity chromatography.

**Fig. 1.**
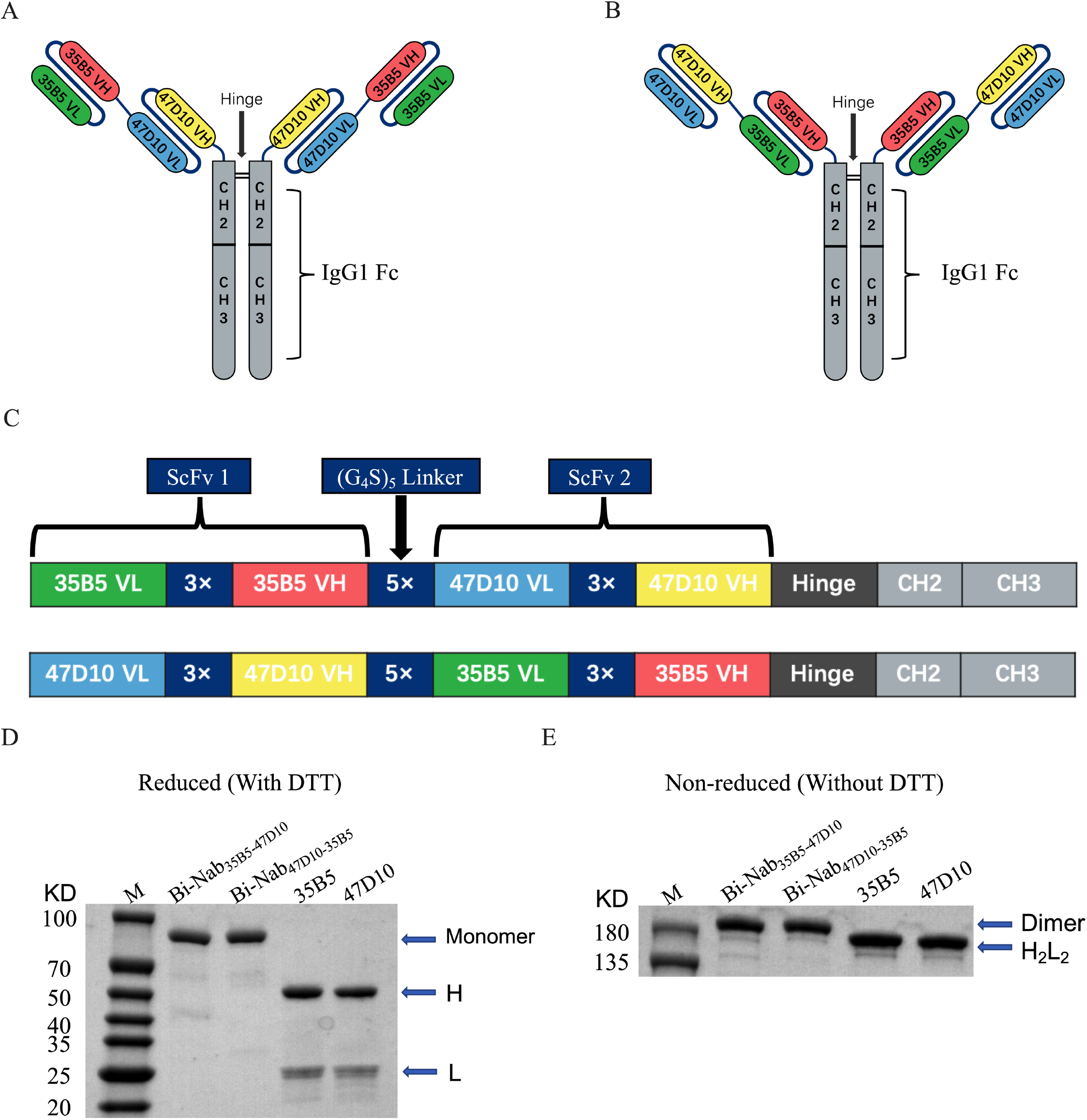
Construction and generation of bsAbs. (**A** and **B**) Overview of the strategy for designing bsAbs. Schematic diagram of the molecular configurations of Bi-Nab_35B5-47D10_ (A) and Bi-Nab_47D10-35B5_ (B). (C) Schematic presentation of the bsAbs. Antibody domains are color-coded according to their architecture (green, variable light chain of 35B5; red, variable heavy chain of 35B5; blue, variable light chain of 47D10; yellow, variable heavy chain of 47D10; gray, human IgG1 Fc). (**D**) Reduced SDS–PAGE analysis of two bsAbs and two parental mAbs. The proteins were analyzed under reducing conditions (+DTT). H and L denote the heavy and light chains, respectively. The molecular weight of the bsAbs monomer was 90 kD. M, molecular weight standard. (**E**) Nonreduced SDS–PAGE analysis of affinity purified bsAbs and parental mAbs. All four antibodies were expressed in ExpiCHO-S^TM^ cells and captured on Protein-A affinity resin. The proteins were analyzed under direct affinity elution conditions (without DTT). Three independent experiments were performed at this scale with the same results. Additional independent experiments yielded similar results at larger culture volumes.

To evaluate the biological activity of Bi-Nab_47D10-35B5_ and Bi-Nab_35B5-47D10_, we initially verified whether they were correctly folded soluble proteins by reducing and nonreducing SDS–PAGE (Fig. 1D and E). As expected, both Bi-Nab_47D10-35B5_ and Bi-Nab_35B5-47D10_ ran as homogeneous species at the expected molecular weight: monomer 90 kD in the reducing SDS–PAGE gel and dimer 180 kD in the nonreducing SDS– PAGE gel (Fig. 1D and E). The results indicate that eukaryotically expressed bsAbs fold correctly.

### Binding and inhibiting properties of bsAbs

Next, to validate the simultaneous spike engagement of the two arms of the bsAbs, we performed an indirect ELISA binding assay. The equal concentrations of S1 protein and S2 protein of the WT SARS-CoV-2 strain were used as the coating antigens (Fig. 2A-B). Bi-Nab_35B5-47D10_ showed the best binding ability to the WT SARS-CoV-2 S1, with the lowest concentration for 50% of maximal effect (EC_50_) values of 4.2 ng/ml (Fig. 2A and 2E). Both Bi-Nab_35B5-47D10_ and Bi-Nab_47D10-35B5_ display similar affinity to the S2 protein, although a reduced affinity of bsAbs to the S2 protein compared with the parental mAb 47D10 was observed (Fig. 2B). These findings confirm that Bi-Nab_47D10-35B5_ and Bi-Nab_35B5-47D10_ can simultaneously bind both the RBD and the S2 epitopes, demonstrating that both arms of the bsAbs are functional. Simultaneously, we assessed the potential of bsAbs to block the binding of angiotensin-converting enzyme 2 (ACE2) to the WT SARS-CoV-2 RBD using ELISA-based binding inhibition assays as previously described [37]. Consistent with the binding results of bsAbs to WT S1, the Bi-Nab_35B5-47D10_ exhibited a similar potent blocking interaction between RBD and ACE2 as parental 35B5 and is superior to Bi-Nab_47D10-35D5_ (Fig. 2C). Furthermore, Bi-Nab_35B5-47D10_ also substantially bound to the mutated S1 proteins of SARS-CoV-2 VOCs, which included HV69-70 deletion and N501Y and D614G mutations, and harbored a similar binding capacity to the VOC S1 with parental 35B5 (Fig. 2D). Indeed, Bi-Nab_35B5-47D10_ demonstrated activity equal to, or in some cases better than, that of parental mAbs (Fig. 2E). These results indicate that Bi-Nab_35B5-47D10_ retains the binding capacity and breadth of its parental mAbs.

**Fig. 2.**
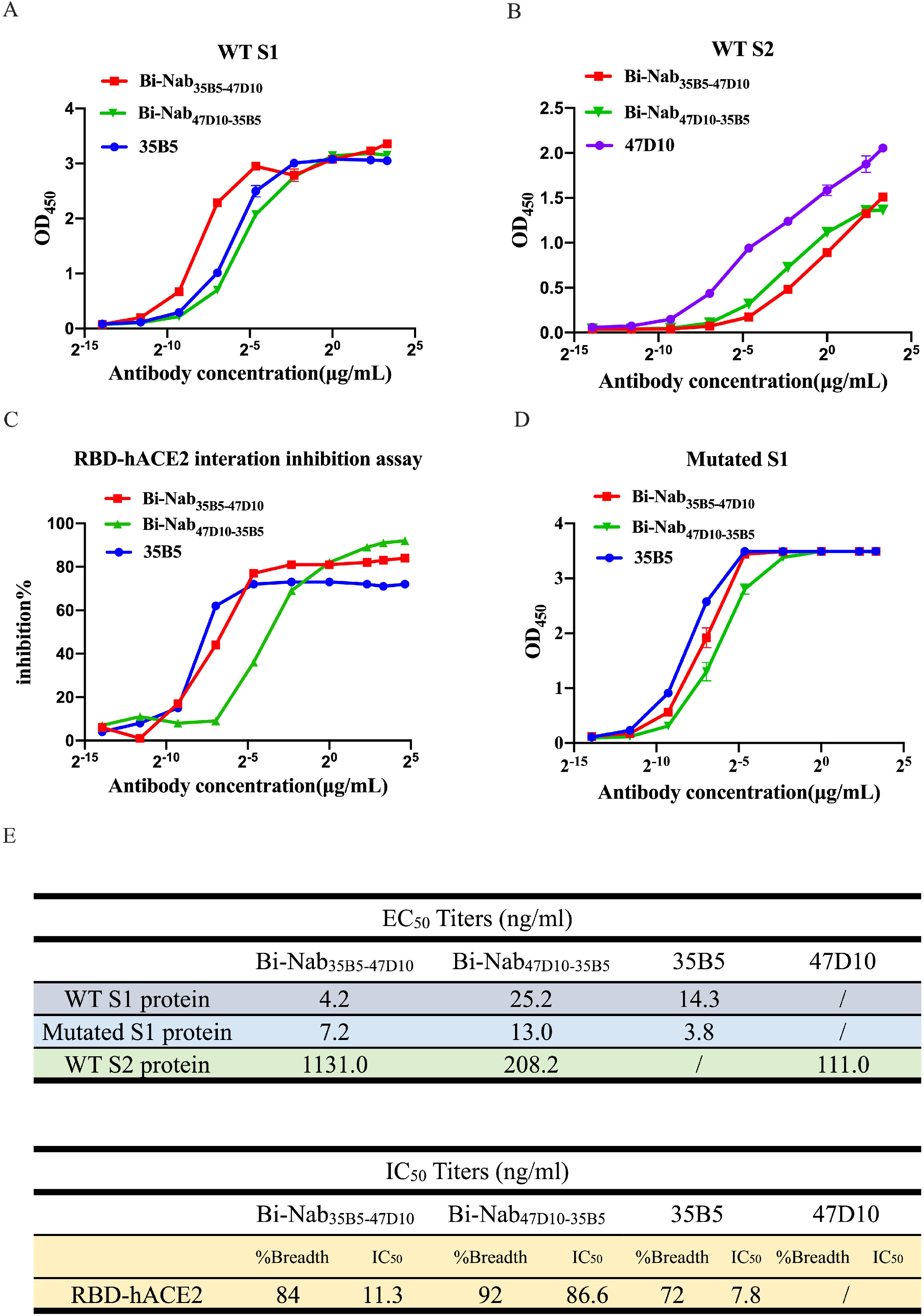
Binding and inhibition properties of bsAbs. (**A** and **B**) ELISA binding assay of bsAbs and parental mAbs to S1 proteins (**A**) or S2 protein (**B**) of WT SARS-CoV-2. EC_50_, concentration for 50% of maximal effect. (**C**) ELISA analysis of bsAbs or parental mAbs-mediated inhibition of WT RBD proteins binding to ACE2. IC_50_, half maximal inhibitory concentration. (**D**) ELISA analysis of bsAbs or parental mAbs binding to the mutated S1 protein of SARS-CoV-2, including HV69-70 deletion, N501Y and D614G. (**E**) Representative EC_50_ and IC_50_ titers (in nanograms per milliliter) of bsAbs and parental mAbs showing the effective binding and inhibiting activity of Bi-Nab_35B5-47D10_.

### Crossing activity of bsAbs on SARS-CoV-2 VOCs and VBM S proteins

Next, we further investigated whether bsAbs could recognize different coronavirus S proteins in the native conformation. Human embryonic kidney (HEK) 293T cells transiently expressing RaTG S protein, SARS-CoV S protein, WT SARS-CoV-2 S protein, Alpha variant (B.1.1.7, N501Y in RBD) S protein [38], Beta variant (B.1.351, K417N, E484K, and N501Y in RBD) S protein [39, 40] or Delta variant (B.1.617.2, L452R, T478K in RBD) S protein [41] were incubated with Bi-Nab_35B5-47D10_, Bi-Nab_47D10-35B5_, mAb 35B5 or mAb 47D10, respectively, followed by flow cytometry analysis. ACE2 was used as a positive control (Fig. 3A). Consistent with the previous results that mAb 35B5 is a SARS-CoV-2-specific neutralizing antibody targeting a conserved epitope on RBD, 35B5 easily detected the expression of four different SARS-CoV-2 S proteins on the HEK293T cell surface with no cross-reactivity of the RaTG S protein (Fig. 3B). Neither two parental mAbs nor two bsAbs showed a binding signal for the SARS-CoV S protein (data not shown). We found that mAb 47D10 cross-reacted with RaTG S protein (Fig. 3C), which expands the minor cross-reaction of Bi-Nab_35B5-47D10_ and Bi-Nab_47D10-35B5_ to RaTG S protein (Fig. 3D and E). The surface spike protein of Alpha variant, Beta variant and Delta variant was readily detected by both Bi-Nab_35B5-47D10_ and Bi-Nab_47D10-35B5_ just as parental 35B5 mAb. Notably, both bsAbs significantly increased the affinity of the Delta variant S protein (Fig. 3D and 3E). Taken together, Bi-Nab_35B5-47D10_ can specifically bind to the native S proteins of SARS-CoV-2 VOCs and VBMs (Fig. 3F).

**Fig. 3.**
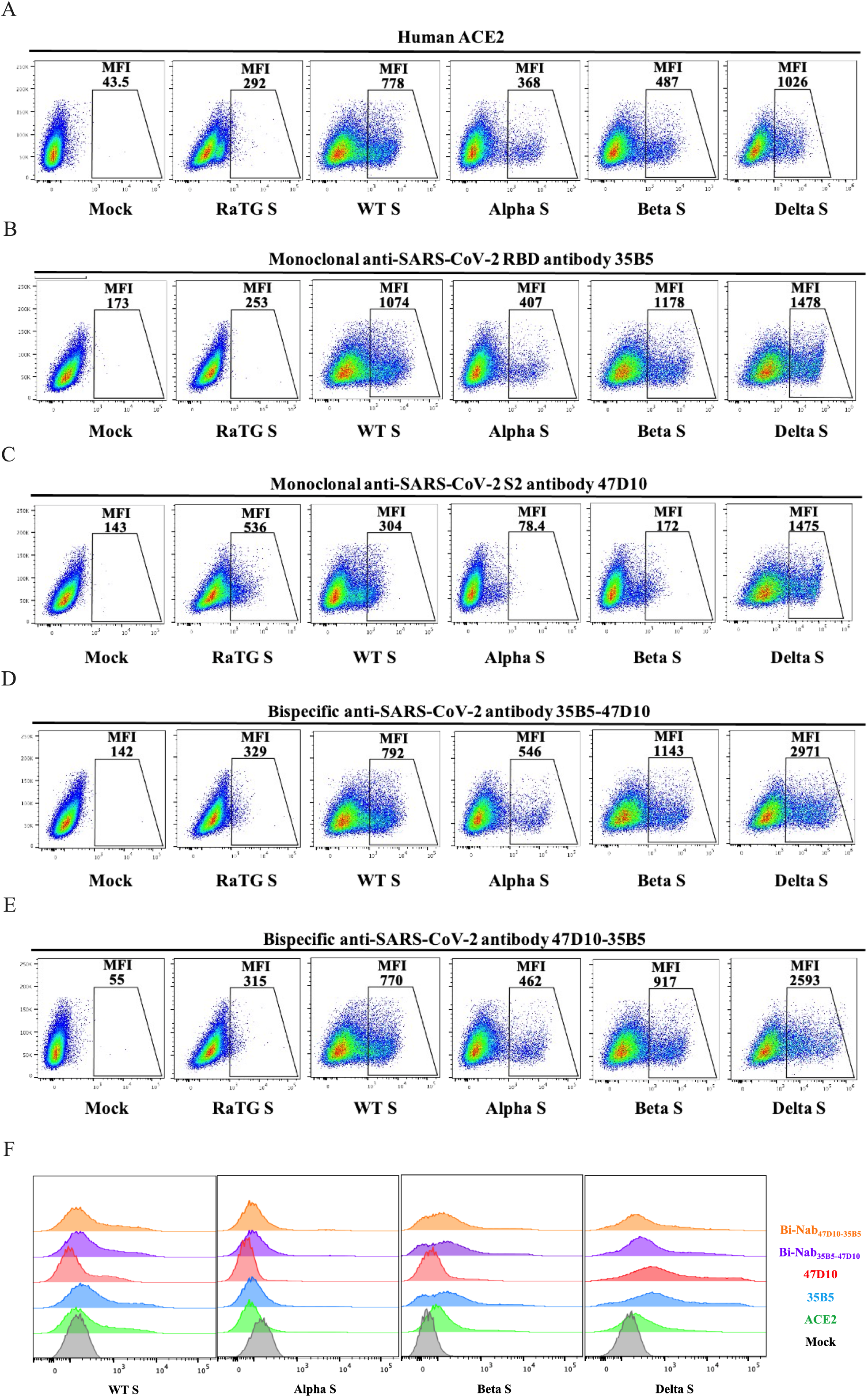
Binding kinetics of bsAbs to cell surface-associated coronavirus S protein. (**A-E**) Binding of bsAbs or parental mAbs to RaTG S, WT SARS-CoV-2 S, Alpha S, Beta S or Delta S proteins. HEK293T cells were transfected to transiently express RaTG S, WT SARS-CoV-2, Alpha S, Beta S or Delta S proteins and incubated with mAb 35B5 (**B**), mAb 47D10 (**C**), Bi-Nab_35B5-47D10_ (**D**) and Bi-Nab_47D10-35B5_ (**E**), respectively, for 1 h on ice. Soluble hACE2 with a His tag was used as a positive control (**A**), followed by a FITC-conjugated anti-human IgG Fc or FITC-conjugated anti-His antibody. Then, the cells were analyzed by flow cytometry. Mean fluorescence intensity (MFI) was normalized to the empty vector (mock) group (**F**). The experiments were performed three times, and one representative is shown.

### Elevated neutralization sensitivity of Bi-Nab_35B5-47D10_ to SARS-CoV-2 VBM variants

The emergence of SARS-CoV-2 VBMs with various mutations including Alpha (with N501Y in RBD, T716I, S982A and D1118H in S2), Beta (with K417N, E484K, and N501Y in RBD, A701 V in S2), Gamma (with K417T, E484K and N501Y in RBD, T1027I and V1176F in S2), Epsilon (with L452R in RBD), Eta (with E484K in RBD, F888 L in S2), Iota (with S477N and E484K in RBD, A701 V in S2), Mu (with R346K, E484K and N501Y in RBD, D950N in S2), Zeta (with E484K in RBD, V1176F in S2), 1.617.3 (with L452R and E484K in RBD, D950N in S2) and Kappa (with L452R and E484Q in RBD, Q1071H in S2) escape neutralization and threaten efforts to contain the COVID-19 pandemic (Fig. 4A).

**Fig. 4.**
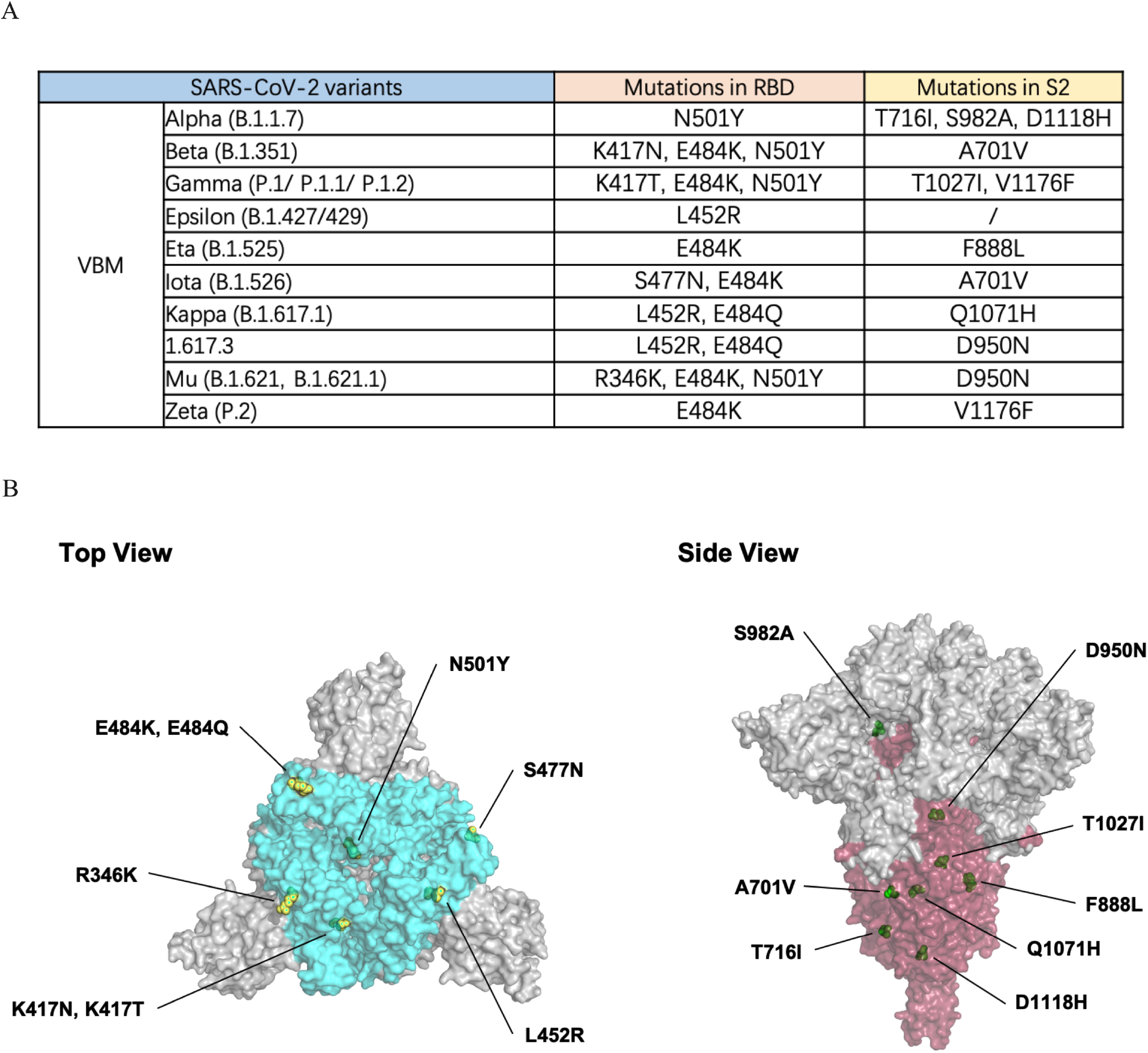
RBD and S2 mutations of VBM variants. (**A**) Statistics on VBM RBD and S2 mutations are displayed. (**B**) The crystal structure of the SARS-CoV-2 spike trimer (PDB ID 7KRQ) highlighting the mutational landscape of VBM variants relative to WT SARS-CoV-2. The epitopes of the RBD (bright blue) and S2 (dark red) regions are shown. The mutations are indicated by yellow (RBD mutations) and green (S2 mutations) spheres on the surface of the S trimer using the PyMOL software suite.

The rational design of bsAb is to avoid escape mutants. To initially assess the virus neutralization capacity of the two bsAbs in comparison to their parental mAbs 35B5 and 47D10, we performed neutralization assays with pseudovirus. Because the two parental mAbs target the RBD and S2, respectively, we focused on mutations of VBMs that occur in the RBD and S2 regions (Fig. 4B). Alpha, beta and kappa variants were chosen to represent VBMs because these variants contain most of the VBM mutations (Fig. 4A).

WT SARS-CoV-2, alpha, beta and kappa variants S protein pseudovirus were incubated with 35B5, 47D10, Bi-Nab_35B5-47D10_ or Bi-Nab_47D10-35B5_, respectively, and their transduction was measured according to luciferase activities. As expected, Bi-Nab_35B5-47D10_ neutralized WT SARS-CoV-2 S pseudovirus effectively with half maximum inhibitory concentration (IC_50_) values of 8.2 ng/ml (Fig. 5A) and displayed substantially improved neutralization breadth among the three VBM variants, especially for alpha and kappa variants (Fig. 5B-D). Among the four antibodies, Bi-Nab_35B5-47D10_ and Bi-Nab_47D10-35B5_ potently neutralized Alpha and Kappa variants and exhibited comparable neutralization potencies with much lower IC_50_ values than parental mAbs (Fig. 5B, 5D and 5E). Among the three VBM variants, the beta variant exhibited less sensitivity to the 35B5 mAb (Fig. 5C). Notably, Bi-Nab_35B5-47D10_ increased the neutralization activity against the beta pseudovirus approximately 6-fold, with the lowest IC_50_ value of 64.5 ng/ml compared with 35B5 (Fig. 5C and 5E). Therefore, these results suggest enhanced cross-reactivity of Bi-Nab_35B5-47D10_, as indicated by the elevated potency against the WT SARS-CoV-2, Alpha, Beta and Kappa variants (Fig. 5E).

**Fig. 5.**
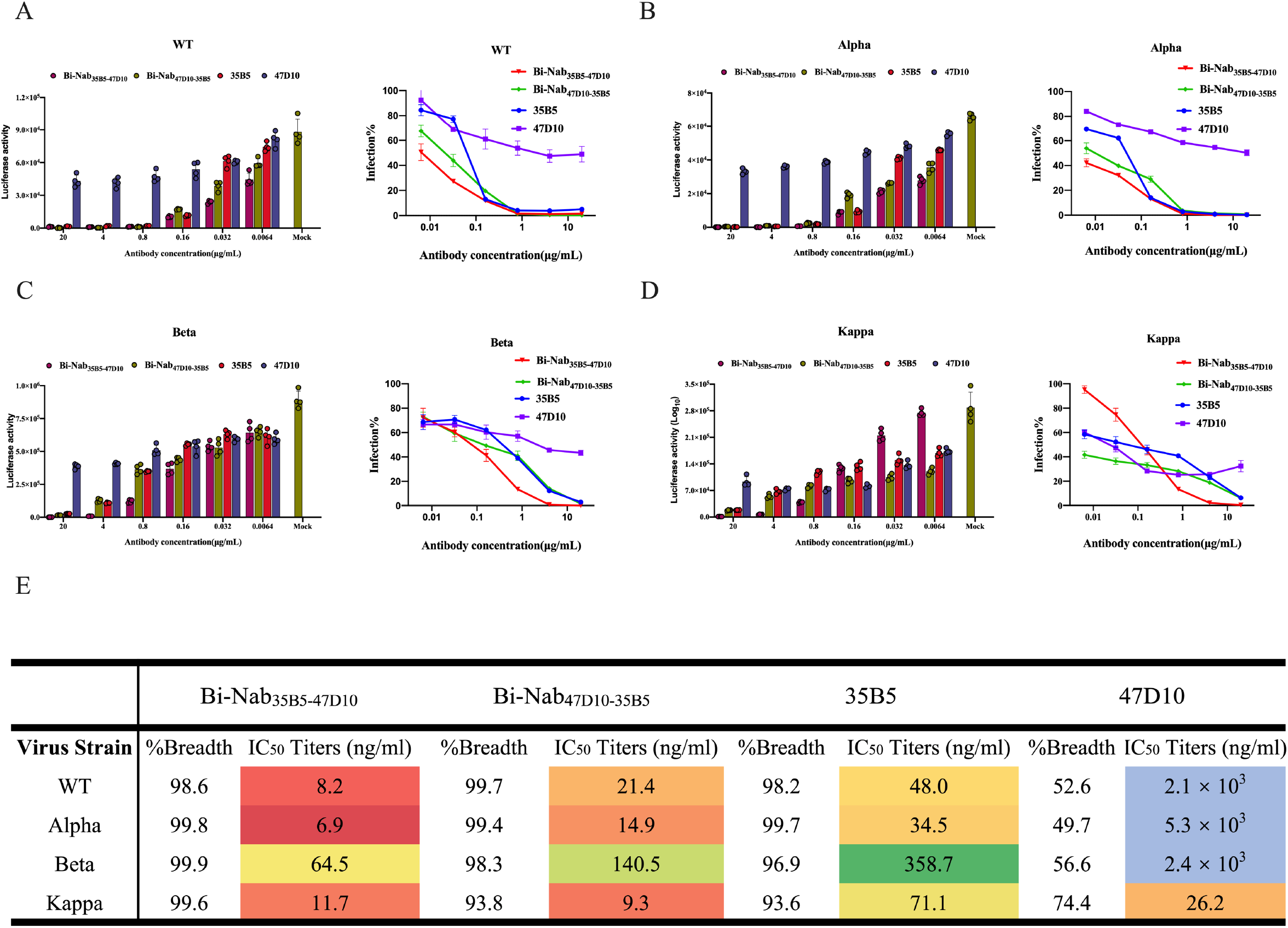
Neutralization of bsAbs against WT and VBM pseudoviruses. (**A-D**) Two parental mAbs and two bsAbs mediated neutralization of the indicated pseudovirus. WT SARS-CoV-2 S (**A**), Alpha S (**B**), Beta S (**C**) or Kappa S (**D**) pseudovirus were preincubated with fivefold serially diluted Bi-Nab_47D10-35B5_, Bi-Nab_35B5-47D10_, 47D10 or 35B5. Then, the mixture was added to HEK293 cells transiently expressing hACE2 and lysed 48 h later, and their transduction was measured according to luciferase activities. Potencies were calculated against sensitive viruses, and heatmaps of IC_50_ titers were generated in Excel. Warmer colors indicate more potent neutralization. Breadths based on IC_50_s are also summarized (E). Representative IC_50_ titers (in nanograms per milliliter) and neutralization breadth of bsAbs and the parental mAbs showing the improved neutralization activity of Bi-Nab_35B5-47D10_. The experiment was performed twice, and a representative is shown. Error bars represent the SEM of technical triplicates.

### Bi-Nab_35B5-47D10_ potently neutralizes SARS-CoV-2 Delta and Omicron variants

The delta variant and omicron variant are more transmissible than other variants and are linked to a resurgence of COVID-19 in many countries across the world. Finally, we set out to identify the neutralization properties of bsAbs on SARS-CoV-2 VOC delta and submicron variants. Compared to the mutations found in Delta, the Omicron lineage harbors more than thirty amino acid mutations in the spike protein, and most mutations are structurally focused in RBD (Fig. 6A) in regions accessible to antibodies at the top of the spike, increasing the likelihood of immune evasion (Fig. 6B) [42]. Parental mAbs 35B5 and 47D10 exhibited comparable neutralization potencies against the Delta variant, with IC_50_ values of 54.9 ng/ml and 45.2 ng/ml (Fig. 7A), respectively, whereas the neutralizing activity of mAbs 35B5 and 47D10 against Omicron variants was reduced substantially, possibly due to the more mutations in both S1 and S2 regions than Delta (Fig. 6B). 47D10 only slightly neutralized Omicron BA.1 with an IC_50_ value of 1661.0 ng/mL (Fig. 7B); at the same time, it did not effectively neutralize Omicron BA.2 (Fig. 7C). These results are consistent with published studies showing that Omicron displays enhanced neutralization escape compared with other SARS-CoV-2 variants. Interestingly, although Bi-Nab_47D10-35B5_ exhibited substantially lower but detectable neutralization (146.1 ng/ml for Delta, 273.7 ng/ml for Omicron BA.1, 519.2 ng/ml for Omicron BA.2). In comparison with 35B5, Bi-Nab_35B5-47D10_ showed substantially higher neutralization titers against Delta, Omicron BA.1and Omicron BA.2 variants than Bi-Nab_47D10-35B5_ and parental mAbs with IC_50_ values of 14.3 ng/ml, 27.6 ng/ml and 121.1 ng/ml, respectively (Fig. 7D). This indicates that the molecular topology of bsAbs substantially impacts the neutralizing activity against VOC variants. Notably, the neutralizing activity of Bi-Nab_35B5-47D10_ against Omicron was largely enhanced when linking the scFv of 35B5 and 47D10, although mAb 47D10 only slightly neutralized Omicron (Fig. 7D). Collectively, Bi-Nab_35B5-47D10_ potently neutralized both the Delta and Omicron variants and substantially increased the neutralization activity in comparison with their parental mAbs.

**Fig. 6.**
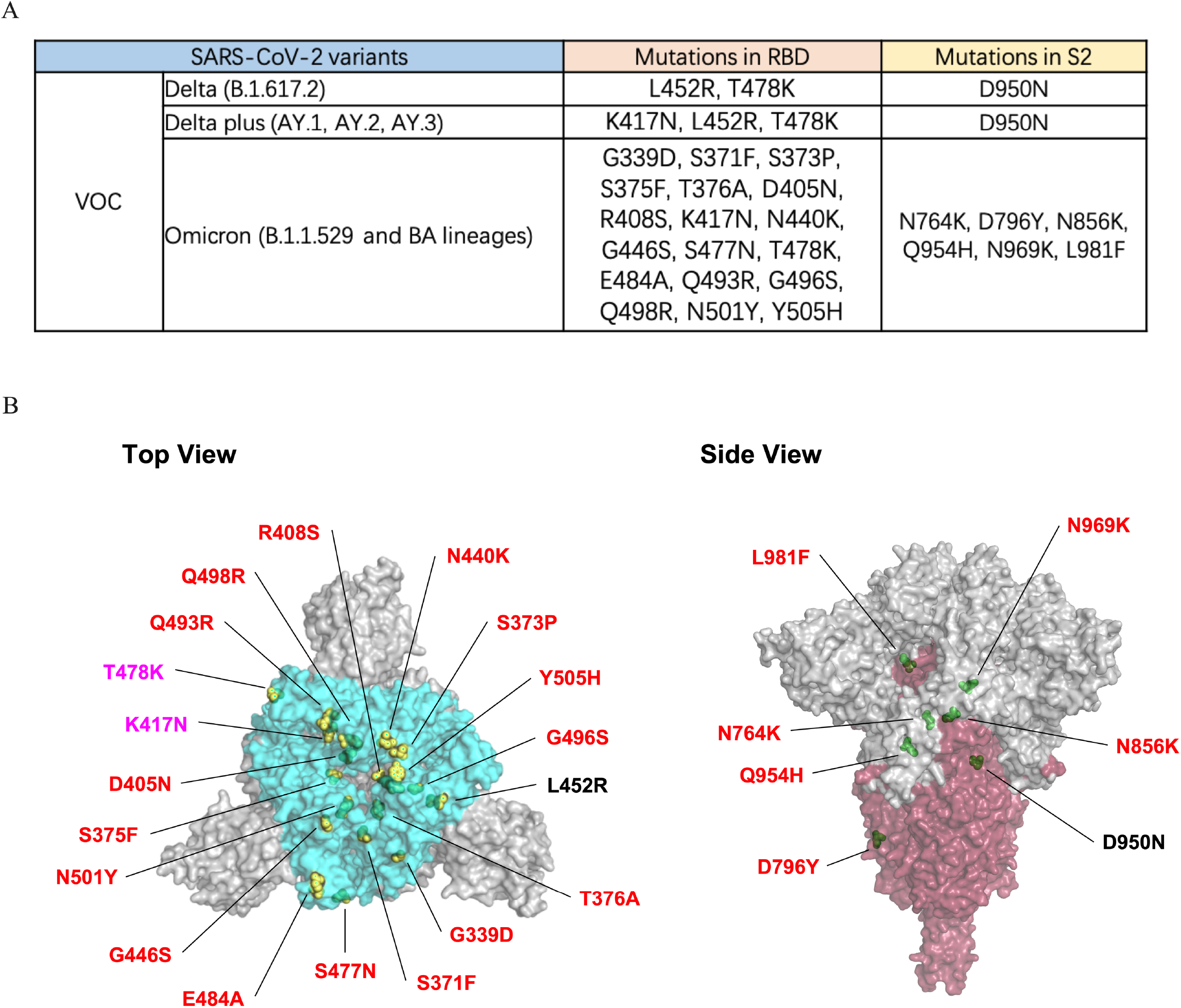
RBD and S2 mutations of VOC variants. (**A**) Schematic of VOC RBD and S2 mutations is illustrated. (**B**) Top view (left panel) and side view (right panel) of spike trimer are shown with mutations in RBD and S2 and highlighted with residue atoms as colored spheres indicated in yellow (RBD mutations, the red font indicates the mutations unique to Omicron variant and the pink refers to the mutations common to VOCs) and green (S2 mutations) on the surface of the S trimer (PDB ID 7KRQ).

**Fig. 7.**
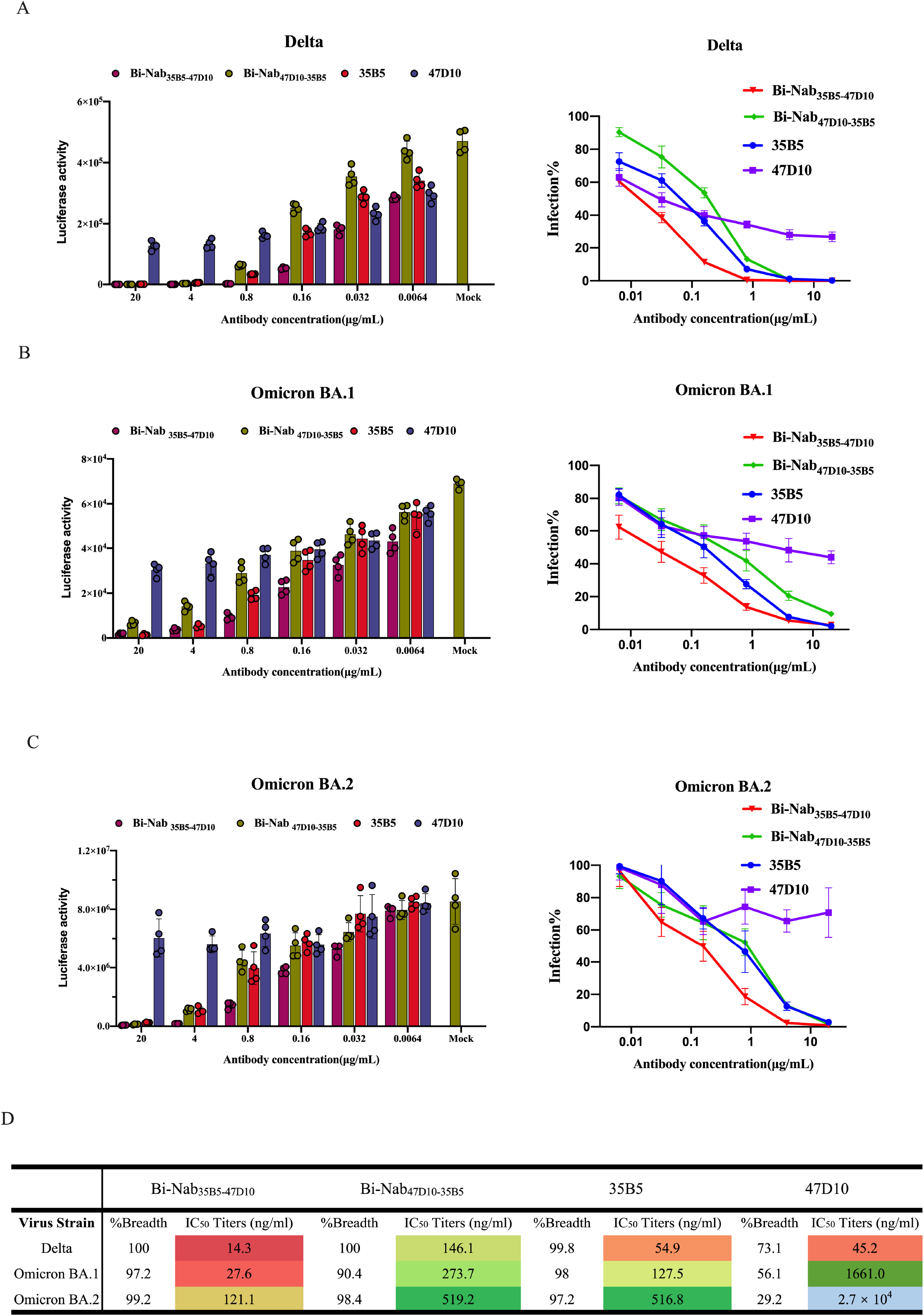
Cross-reactive neutralization of bsAbs against delta and submicron pseudoviruss. Using a lentiviral-based pseudovirus system, the neutralization potency of two parental mAbs and two bsAbs against Delta (**A**), Omicron BA.1 (**B**) and Omicron BA.2 (**C**) pseudoviruses were analyzed. The IC_50_ was determined by log (inhibitor) response of nonlinear regression, and bars and error bars depict the mean and standard error of the mean. (**D**) Bi-Nab_35B5-47D10_ exhibited significantly improved neutralization activity compared to Bi-Nab_47D10-35B5_ and parental mAbs. IC_50_ titers (in nanograms per milliliter), breadth and potency of two bsAbs and two parental mAbs against Delta and Omicron pseudovirus are presented with heatmaps.

## Discussion

The ongoing evolution of SARS-CoV-2 and the emergence of new SARS-CoV-2 variants compromise the efficacy of current SARS-CoV-2 vaccines and licensed mAb therapies[43, 44]. The unusually high mutations of the Omicron variant with high spreads have resulted in breakthrough infections in the world since its first report in November 2021[45, 46]. It is still urgent to develop highly potent and broadly neutralizing mAbs targeting multiple SARS-CoV-2 variants[9, 47]. In this study, we generated a potent bispecific mAb Bi-Nab_35B5-47D10_ targeting spike RBD and S2 in two distinct regions. Our results demonstrate that Bi-Nab_35B5-47D10_ retained the specificity of their parental mAbs and increased potency and breadth compared with their parental mAbs. Bi-Nab_35B5-47D10_ exhibited pan-neutralizing activities against WHO-stated SARS-CoV-2 VBMs and VOCs, including B.1.617.2 (Delta), Omicron BA.1 and Omicron BA.2 variants, highlighting its potential application in the prevention and treatment of SARS-CoV-2 VOCs.

Bispecific or multiple specific antibodies targeting different regions of the spike protein are a favorable strategy to treat COVID-19 caused by the ongoing emergence of new SARS-CoV-2 VOCs[48, 49]. Except for the increased threshold to produce neutralizing escape mutants, bispecific antibodies have practical and cost advantages over the mAb cocktail strategy since the complicated formulation of mAb cocktails routinely increases manufacturing costs and volumes. Here, we explored two SARS-CoV-2 neutralizing mAbs, 35B5 and 47D10, which target divergent spike regions and block SARS-CoV-2 infection by distinct mechanisms. 35B5 binds to an invariant epitope in the RBD and causes dissociation of the spike trimer by a glycan displacement action[35]. The epitope of 35B5 in the RBD is invariant in SARS-CoV-2 WT, four VBMs and two Delta and Omicron VOCs[35, 36]. In addition to the earlier Omicron variants BA.1 and BA.2, more Omicron subvariants have emerged, including BA.3, BA.4 and BA.5[50, 51]. Among the identified Omicron subvariants, the BA.2 subvariant is more contagious than other subvariants and is now the dominant strain globally[52]. Fortunately, the epitope of 35B5 on RBD does not contain the mutations of Omicron BA.1 and BA.2 (Fig. 6)[35, 50]. Crucially, Bi-Nab_35B5-47D10_ further improved the potency to neutralize Omicron VOC in comparison with parental 35B5 (Fig. 7). Therefore, it is rational that Bi-Nab_35B10-47D10 will_ potently be of interest for further clinical development for COVID-19 treatment.

## Materials and Methods

### Design and expression of bispecific antibodies

Plasmids containing the heavy and light chain genes for the production of the monoclonal antibodies 35B5 and 47D10 were prepared as previously described [53]. Single-chain Fv format bispecific antibodies were designed from the sequences of the variable regions of monoclonal antibodies 35B5 and 47D10 (ScFv 35B5-47D10) or 47D10 and 35B5 (ScFv 47D10-35B5), utilizing tandem glycine-serine (G4S) peptide linkers. Codon-optimized ScFvs DNA sequences were synthesized and cloned into the pUC57 cloning vector (GenScript, Piscataway, NJ) and subcloned into the eukaryotic cell expression vector AbVec-hIgG1 between the AgeI and Hind III sites.

The ExpiCHO^TM^ Expression System (Thermo Fisher) was used to produce bsAbs. Briefly, following the manufacturer’s max titer protocol, 25 ml (6×10^6^ cells/ml) ExpiCHO-S^TM^ cells in a 125 ml flask were transfected with a master mixture containing 25 μg bispecific antibody plasmid and 80 μl ExpiFectamine^TM^ CHO reagent diluted in 1 ml cold OptiPRO^TM^ SFM complexation medium. The ExpiFectamine^TM^ CHO/DNA complexes were added to the cells immediately or after up to 5 minutes of incubation at room temperature without any loss of performance. The cells were incubated on an orbital shaker at 37°C with a humidified atmosphere of 8% CO_2_ humidified in air without the need to change or add media. On the day after transfection, 150 μl ExpiCHO^TM^ enhancer and 6 ml ExpiCHO^TM^ feed were added to the flask. A second volume of ExpiCHO^TM^ feed was added to cultured cells on day 5 posttransfection, and the flask was immediately returned to the shaking incubator. Supernatants were harvested at 4000-5000 x g for 30 minutes in a refrigerated centrifuge, and the supernatant was filtered through a 0.22 μm filter on day 12 posttransfection.

### Bispecific antibody isolation and purification

Briefly, bsAbs were efficiently purified by using Protein A Sepharose affinity chromatography medium (GenScript, L00210-10). The purified antibodies were separated on a 7.5%–12% polyacrylamide gel and revealed with Coomassie blue under reducing or nonreducing conditions. To assure functionality, stability, and batch-to-batch consistency, all antibodies were subjected to quality control and biophysical characterization.

### Cells and plasmids

HEK293T cells producing pseudovirus and HEK293 cells overexpressing recombinant human ACE2 (293/hACE2) were preserved in our laboratory and maintained in Dulbecco’s modified Eagle’s medium (DMEM, Thermo Fisher) containing 10% fetal bovine serum (FBS, Gibco) and 1% penicillin–streptomycin and were incubated at 37ºC with 5% CO_2_ and 95% humidity. ExpiCHO-S^TM^ cells (Thermo Fisher) were cultured in ExpiCHO^TM^ expression medium in 125 mL shaker flasks in a 37°C incubator with ≥ 80% relative humidity and 8% CO_2_ on an orbital shaker with a 50 mm shaking diameter rotating at 95 rpm. For routine maintenance, ExpiCHO-S^TM^ cells were typically passaged every 3 days at a ratio of 1:20 when they were grown to 3-5×10^6^ cells/mL. The pcDNA3.1-hACE2 plasmid with human codon optimization, plasmids encoding WT SARS-CoV-2 S glycoprotein, SARS-CoV-2 VBM spike protein, SARS-CoV-2 VOC S glycoprotein, RaTG S glycoprotein, lentiviral packaging plasmid psPAX2 and pLenti-GFP lentiviral reporter plasmid were generously gifted by Dr. Zhaohui Qian.

### Enzyme-linked immunosorbent assay

To evaluate antibody characterization in vitro, the ELISA method with modifications was used as reported previously[54]. In brief, 50 nanograms (ng) of SARS-CoV-2 S1 protein of the WT strain (Sino Biological, 40591-V08H) or B.1.1.7 (Sino Biological, 40591-V08H7) or SARS-CoV-2 S2 (Sino Biological, 40590-V08B) protein or SARS-CoV-2 RBD protein (Sino Biological, 40592-V08B) was coated on ELISA plates in 100 μl per well at 37°C for 2 hours or 4°C overnight. After washing 3 times with PBST, blocking buffer with 5% FBS and 0.05% Tween 20 was added to the ELISA plates and incubated for 1 hour at 37°C. Next, 100 mL serially diluted mAbs or bsAbs was added to each well in 100 μl blocking buffer for 1 hour at 37°C. Following washing 3 times, mouse anti-human IgG Fc secondary antibody with HRP (Abcam) was added and incubated at 37°C for 1 h, followed by washing with PBST. The ELISA plates were reacted with 3,30,5,50-tetramethylbenzidine (TMB, Sigma) substrate at 25°C for 5 minutes and then stopped by 0.2 M H_2_SO_4_ stop buffer. The optical density (OD) at 450 nm was measured using an iMark microplate absorbance reader (BIO-RAD). Nonlinear regression was used to calculate the EC_50_.

### ELISA-based receptor-binding inhibition of hACE2

The ability of antibodies to inhibit the binding of the SARS-CoV-2 RBD to hACE2 was investigated by ELISA. The 96-well ELISA plates were coated with 200 ng of hACE2 protein (Sino Biological, 10108-H08H) per well overnight at 4°C, washed with PBST and blocked for 1 hour with blocking buffer as above. Meanwhile, fivefold serial dilutions of mAbs or bispecific antibodies were incubated with 4 ng/mL SARS-CoV-2 RBD with mouse IgG FC tag protein (Sino Biological, 40592-V05H) for 1 hour at 25°C. Then, the mixtures were added to ELISA plates and incubated for 1 hour at 37°C. After further washing, bound SARS-CoV-2 RBD protein was detected with anti-mouse Fc HRP antibody (Abcam) diluted 1:10000 in blocking solution followed by washing with PBST. The ELISA plates were reacted with TMB substrate at 25°C for 5 minutes and then stopped by 0.2 M H_2_SO_4_ stop buffer and determined at OD 450 nm. The IC_50_ was determined by using 4-parameter logistic regression.

### SARS-CoV-2 pseudotyped reporter virus and pseudotyped virus neutralization assay

Generation of SARS-CoV-2 or SARS-CoV-2 VOC-type pseudoviruss was performed as previously described[55]. In brief, pseudoviruses were produced by using PEI to cotransfect 293T cells with psPAX2, pLenti-GFP and plasmids encoding SARS-CoV-2 S, SARS-CoV-2 VOCs S, SARS-CoV S, RaTG S or empty vector at a ratio of 1:1:1. The media was replaced with fresh media containing 10% fetal bovine serum and 1% penicillin–streptomycin 4 hours post-transfection. The supernatants were harvested 48 hours post-transfection and centrifuged at 800 ×g for 5 min to remove cell debris before passing through a 0.45 μm filter.

For the pseudovirus neutralization assay, HEK293 (hACE2/293) cells 80-90% confluent in T75 cell culture flasks were transfected with 20 μg of plasmid encoding hACE2 for 36 hours and seeded into 24-well plates the day before transduction with pseudovirus. Fivefold serially diluted bsAbs or mAbs were incubated with SARS-CoV-2 pseudotyped virus for 1 hour. The 500 μl per well mixture was subsequently incubated with hACE2/293 cells overnight, and then the mixture was changed to fresh media. Approximately 48 hours post-incubation, the luciferase activity of SARS-CoV-2-type pseudovirus-infected hACE2/293 cells was detected by the Dual-Luciferase Reporter Assay System (Promega), and cells were lysed with 120 μl medium containing 50% Steady-Glo and 50% complete cell growth medium at room temperature for 5 minutes. The percentage of infection was normalized to those derived from cells infected with SARS-CoV-2 pseudotyped virus in the absence of antibodies. The IC_50_ was determined by using 4-parameter logistic regression.

### Flow cytometry-based bsAb binding assay

HEK293T cells 80-90% confluent in 6-well cell culture plates were transfected with 2 μg of plasmids encoding either WT SARS-CoV-2 S or SARS-CoV-2 mutated S protein or SARS-CoV S protein or RaTG S protein using linear polyethylenimine (PEI, Sigma–Aldrich, 408727). A total amount of 2 μg DNA diluted in 1 ml per well of DMEM was mixed with PEI in a 1:2 ratio. The transfection mixture was added to cell culture plates with DMEM in a dropwise manner after 15 minutes of incubation at room temperature. After 4-6 hours of incubation at 37°C under 5% CO_2_, the medium was changed to fresh complete cell growth medium (DMEM supplemented with 10% FBS and 1% penicillin–streptomycin). Cells were detached by using PBS with 1 mM EDTA 40 hours post-transfection and washed with cold PBS containing 2% FBS. After washing, the cells were incubated with bsAbs, monoclonal human anti-SARS-CoV-2 RBD antibody 35B5 or monoclonal human anti-SARS-CoV-2 S2 antibody 47D10 (5 μg per well) for 1 hour on ice, followed by FITC-conjugated anti-human IgG (1:100) (ZSGB-Bio, ZF-0308) for 1 hour on ice away from light. Cells were acquired by flow cytometry (BD Biosciences) and analyzed using FlowJo.

### Statistical analysis

All statistical analyses were performed using GraphPad Prism 9.0 software. In the ELISA, three independent experiments were performed, and the mean values ± SEM and the EC_50_ values were calculated by using sigmoidal dose–response nonlinear regression. In the ELISA-based inhibition assay and pseudovirus neutralization assay, three or two independent experiments were performed, and the mean values ± SEM and the IC_50_ values were plotted by fitting a nonlinear four-parameter dose–response curve. The figure legends show all of the statistical details of the experiments. PyMol was used to prepare all the structure figures (Schrodinger: https://www.schrodinger.com/pymol).

## Acknowledgments

The work was supported by the National Natural Science Foundation of China (32192453) and the Chinese Universities Scientific Fund (2022RC019). The content is solely the authors’ responsibility and does not necessarily represent the official views of the funding resources.

## Figure legends

**Fig. S1. Isolation of SARS-CoV-2 S2 targeted mAb from COVID-19 convalescent patients.**(**A**) Isolation strategy of SARS-CoV-2 S2 mAb 47D10 from COVID-19 convalescent patients. (**B**) Flow cytometry analysis of SARS-CoV-2 S2-specific B cells from the PBMCs of healthy donors and COVID-19 convalescent patients. The numbers adjacent to the outlined area indicate the proportions of SARS-CoV-2 S2-specific B cells in CD19+CD20+IgG+ B cells.

## References

1. Arora, P., et al., Delta variant (B.1.617.2) sublineages do not show increased neutralization resistance. Cell Mol Immunol, 2021. 18(11): p. 2557–2559.

2. Bruel, T., et al., Serum neutralization of SARS-CoV-2 Omicron sublineages BA.1 and BA.2 in patients receiving monoclonal antibodies. Nat Med, 2022.

3. Mlcochova, P., et al., SARS-CoV-2 B.1.617.2 Delta variant replication and immune evasion. Nature, 2021. 599(7883): p. 114–119.

4. Liu, L., et al., Striking antibody evasion manifested by the Omicron variant of SARS-CoV-2. Nature, 2022. 602(7898): p. 676–681.

5. Planas, D., et al., Reduced sensitivity of SARS-CoV-2 variant Delta to antibody neutralization. Nature, 2021. 596(7871): p. 276–280.

6. Zhou, D., et al., Evidence of escape of SARS-CoV-2 variant B.1.351 from natural and vaccine-induced sera. Cell, 2021. 184(9): p. 2348–2361.e6.

7. Yuan, M., et al., Structural basis of a shared antibody response to SARS-CoV-2. Science, 2020. 369(6507): p. 1119–1123.

8. Cerutti, G., et al., Potent SARS-CoV-2 neutralizing antibodies directed against spike N-terminal domain target a single supersite. Cell Host Microbe, 2021. 29(5): p. 819–833.e7.

9. Barnes, C.O., et al., SARS-CoV-2 neutralizing antibody structures inform therapeutic strategies. Nature, 2020. 588(7839): p. 682–687.

10. McCallum, M., et al., N-terminal domain antigenic mapping reveals a site of vulnerability for SARS-CoV-2. Cell, 2021. 184(9): p. 2332–2347.e16.

11. Wibmer, C.K., et al., SARS-CoV-2 501Y.V2 escapes neutralization by South African COVID-19 donor plasma. Nat Med, 2021. 27(4): p. 622–625.

12. Garcia-Beltran, W.F., et al., Multiple SARS-CoV-2 variants escape neutralization by vaccine-induced humoral immunity. Cell, 2021. 184(9): p. 2372–2383.e9.

13. Odak, I., et al., Longitudinal Tracking of Immune Responses in COVID-19 Convalescents Reveals Absence of Neutralization Activity Against Omicron and Staggered Impairment to Other SARS-CoV-2 Variants of Concern. Front Immunol, 2022. 13: p. 863039.

14. Ye, G., B. Liu, and F. Li, Cryo-EM structure of a SARS-CoV-2 omicron spike protein ectodomain. Nat Commun, 2022. 13(1): p. 1214.

15. Torjesen, I., Covid-19: Omicron may be more transmissible than other variants and partly resistant to existing vaccines, scientists fear. Bmj, 2021. 375: p. n2943.

16. Pulliam, J.R.C., et al., Increased risk of SARS-CoV-2 reinfection associated with emergence of Omicron in South Africa. Science, 2022: p. eabn4947.

17. Carreno, J.M., et al., Activity of convalescent and vaccine serum against SARS-CoV-2 Omicron. Nature, 2022. 602(7898): p. 682–688.

18. Iketani, S., et al., Antibody evasion properties of SARS-CoV-2 Omicron sublineages. Nature, 2022.

19. Cao, Y., et al., Omicron escapes the majority of existing SARS-CoV-2 neutralizing antibodies. Nature, 2022. 602(7898): p. 657–663.

20. Hoffmann, M., et al., The Omicron variant is highly resistant against antibody-mediated neutralization: Implications for control of the COVID-19 pandemic. Cell, 2022. 185(3): p. 447–456.e11.

21. Walter, J.D., et al., Biparatopic sybodies neutralize SARS-CoV-2 variants of concern and mitigate drug resistance. EMBO Rep, 2022. 23(4): p. e54199.

22. De Gasparo, R., et al., Bispecific IgG neutralizes SARS-CoV-2 variants and prevents escape in mice. Nature, 2021. 593(7859): p. 424–428.

23. Huang, Y., et al., Engineered Bispecific Antibodies with Exquisite HIV-1-Neutralizing Activity. Cell, 2016. 165(7): p. 1621–1631.

24. Zhao, Q., Bispecific Antibodies for Autoimmune and Inflammatory Diseases: Clinical Progress to Date. BioDrugs, 2020. 34(2): p. 111–119.

25. Wang, J., et al., A Human Bi-specific Antibody against Zika Virus with High Therapeutic Potential. Cell, 2017. 171(1): p. 229–241.e15.

26. Bournazos, S., et al., Bispecific Anti-HIV-1 Antibodies with Enhanced Breadth and Potency. Cell, 2016. 165(7): p. 1609–1620.

27. Yao, H., et al., Rational development of a human antibody cocktail that deploys multiple functions to confer Pan-SARS-CoVs protection. Cell Res, 2021. 31(1): p. 25–36.

28. Weinreich, D.M., et al., REGN-COV2, a Neutralizing Antibody Cocktail, in Outpatients with Covid-19. N Engl J Med, 2021. 384(3): p. 238–251.

29. Shima, M., et al., Factor VIII-Mimetic Function of Humanized Bispecific Antibody in Hemophilia A. N Engl J Med, 2016. 374(21): p. 2044–53.

30. Suurs, F.V., et al., A review of bispecific antibodies and antibody constructs in oncology and clinical challenges. Pharmacol Ther, 2019. 201: p. 103–119.

31. Hoffmann, M., H. Kleine-Weber, and S. Pöhlmann, A Multibasic Cleavage Site in the Spike Protein of SARS-CoV-2 Is Essential for Infection of Human Lung Cells. Mol Cell, 2020. 78(4): p. 779–784.e5.

32. Walls, A.C., et al., Structure, Function, and Antigenicity of the SARS-CoV-2 Spike Glycoprotein. Cell, 2020. 181(2): p. 281–292.e6.

33. Zhou, P., et al., Broadly neutralizing anti-S2 antibodies protect against all three human betacoronaviruses that cause severe disease. bioRxiv, 2022.

34. Steinhardt, J.J., et al., Rational design of a trispecific antibody targeting the HIV-1 Env with elevated anti-viral activity. Nat Commun, 2018. 9(1): p. 877.

35. Wang, X., et al., 35B5 antibody potently neutralizes SARS-CoV-2 Omicron by disrupting the N-glycan switch via a conserved Spike epitope. Cell Host & Microbe, 2022.

36. Wang, X., et al., A potent human monoclonal antibody with pan-neutralizing activities directly dislocates S trimer of SARS-CoV-2 through binding both up and down forms of RBD. Signal Transduct Target Ther, 2022. 7(1): p. 114.

37. Chen, X., et al., Disease severity dictates SARS-CoV-2-specific neutralizing antibody responses in COVID-19. Signal Transduct Target Ther, 2020. 5(1): p. 180.

38. Brown, K.A., et al., S-Gene Target Failure as a Marker of Variant B.1.1.7 Among SARS-CoV-2 Isolates in the Greater Toronto Area, December 2020 to March 2021. Jama, 2021. 325(20): p. 2115–2116.

39. Hoffmann, M., et al., SARS-CoV-2 variants B.1.351 and P.1 escape from neutralizing antibodies. Cell, 2021. 184(9): p. 2384–2393.e12.

40. Tegally, H., et al., Detection of a SARS-CoV-2 variant of concern in South Africa. Nature, 2021. 592(7854): p. 438–443.

41. Edara, V.V., et al., Infection and vaccine-induced neutralizing antibody responses to the SARS-CoV-2 B.1.617.1 variant. bioRxiv, 2021.

42. Garcia-Beltran, W.F., et al., mRNA-based COVID-19 vaccine boosters induce neutralizing immunity against SARS-CoV-2 Omicron variant. Cell, 2022. 185(3): p. 457–466.e4.

43. Mileto, D., et al., Reduced neutralization of SARS-CoV-2 Omicron variant by BNT162b2 vaccinees’ sera: a preliminary evaluation. Emerg Microbes Infect, 2022. 11(1): p. 790–792.

44. Dejnirattisai, W., et al., Cross-reacting antibodies enhance dengue virus infection in humans. Science, 2010. 328(5979): p. 745–8.

45. He, X., et al., SARS-CoV-2 Omicron variant: Characteristics and prevention. MedComm (2020), 2021. 2(4): p. 838–45.

46. Gobeil, S.M., et al., Structural diversity of the SARS-CoV-2 Omicron spike. Mol Cell, 2022.

47. Zhou, H., et al., Neutralization of SARS-CoV-2 Omicron BA.2 by Therapeutic Monoclonal Antibodies. bioRxiv, 2022.

48. Cho, H., et al., Bispecific antibodies targeting distinct regions of the spike protein potently neutralize SARS-CoV-2 variants of concern. Sci Transl Med, 2021. 13(616): p. eabj5413.

49. Li, Z., et al., An engineered bispecific human monoclonal antibody against SARS-CoV-2. Nat Immunol, 2022. 23(3): p. 423–430.

50. Arora, P., et al., Comparable neutralisation evasion of SARS-CoV-2 omicron subvariants BA.1, BA.2, and BA.3. Lancet Infect Dis, 2022.

51. Khan, K., et al., Omicron sub-lineages BA.4/BA.5 escape BA.1 infection elicited neutralizing immunity. medRxiv, 2022: p. 2022.04.29.22274477.

52. Mahase, E., Omicron sub-lineage BA.2 may have “substantial growth advantage,” UKHSA reports. BMJ, 2022. 376: p. o263.

53. Chen, X., et al., Human monoclonal antibodies block the binding of SARS-CoV-2 spike protein to angiotensin converting enzyme 2 receptor. Cell Mol Immunol, 2020. 17(6): p. 647–649.

54. Yue, S., et al., Sensitivity of SARS-CoV-2 Variants to Neutralization by Convalescent Sera and a VH3-30 Monoclonal Antibody. Front Immunol, 2021. 12: p. 751584.

55. Ou, X., et al., Characterization of spike glycoprotein of SARS-CoV-2 on virus entry and its immune cross-reactivity with SARS-CoV. Nat Commun, 2020. 11(1): p. 1620.

